# An enhanced isothermal amplification assay for viral detection

**DOI:** 10.1101/2020.05.28.118059

**Authors:** Jason Qian, Sarah A. Boswell, Christopher Chidley, Zhi-xiang Lu, Mary E. Pettit, Benjamin L. Gaudio, Jesse M. Fajnzylber, Ryan T. Ingram, Rebecca H. Ward, Jonathan Z. Li, Michael Springer

**Affiliations:** Department of Systems Biology, Harvard Medical School, Boston, MA 02115, USA; Laboratory of Systems Pharmacology, Harvard Medical School, Boston, MA 02115, USA; Biological and Biomedical Sciences Program, Harvard Medical School, Boston, MA 02115, USA; Brigham and Women’s Hospital, Harvard Medical School, Boston, MA 02115, USA; Massachusetts Consortium on Pathogen Readiness, Boston, MA 20115, USA

## Abstract

Rapid, inexpensive, robust diagnostics are essential to control the spread of infectious diseases. Current state of the art diagnostics are highly sensitive and specific, but slow, and require expensive equipment. We developed a molecular diagnostic test for SARS-CoV-2, FIND (Fast Isothermal Nucleic acid Detection), based on an enhanced isothermal recombinase polymerase amplification reaction. FIND has a detection limit on patient samples close to that of RT-qPCR, requires minimal instrumentation, and is highly scalable and cheap. It can be performed in high throughput, does not cross-react with other common coronaviruses, avoids bottlenecks caused by the current worldwide shortage of RNA isolation kits, and takes ~45 minutes from sample collection to results. FIND can be adapted to future novel viruses in days once sequence is available.

**One sentence summary:** Sensitive, specific, rapid, scalable, enhanced isothermal amplification method for detecting SARS-CoV-2 from patient samples.

## Main text

SARS-CoV-2 has rapidly spread around the world with serious consequences for human life and the global economy (*1*). In many countries, efforts to contain the virus have been hampered by a lack of adequate testing (*2*). Rapid, inexpensive, and sensitive testing is essential for contact tracing and isolation strategies to be effective (*3*). While numerous different tests exist, the overwhelming global need for testing has led to limitations in both the supplies of reagents, e.g. swabs and purification kits, and instrumentation, e.g. quantitative polymerase chain reaction (qPCR) or ID NOW machines. In most cases, overcoming these limitations would require scaling of supply lines by several orders of magnitude over current production capacities. Therefore, in an effort to avoid overrun health care systems and high death tolls, many countries have resorted to costly lockdowns.

The ability to reopen economies safely depends crucially on the testing capacity available. Efforts to increase testing capacity have included testing from saliva (*4*), using non-standard storage media or dry swabs (*5*), and eliminating the normal RNA purification step from the standard RT-qPCR tests (*6*, *7*). Strategies such as pooling samples followed by detection using traditional or high throughput sequencing approaches have also been proposed as a way to allow significantly more testing at a highly reduced cost (*8*, *9*). In general, such strategies force a tradeoff between throughput and sensitivity.

Isothermal amplification technologies have long held promise to offer highly sensitive detection at high throughput, and to allow for widely distributed testing including at-home/point-of-need (PON) tests (*10*, *11*). However, isothermal amplification is plagued by nonspecific amplification events that require secondary amplification and detection steps. These steps add extra complexity to the reactions, removing many of the benefits of the isothermal amplification approach. Many ongoing efforts aim to circumvent these problems for SARS-CoV-2 detection. Most of the approaches developed so far still require an extraction step and/or two amplification steps to achieve high specificity, or have low sensitivities that give poor concordance with the gold standard RT-qPCR test (*11*).

We set out to determine the underlying reasons for the poor performance of isothermal amplification technologies in viral detection applications. We selected reverse transcription-recombinase polymerase amplification (RT-RPA) as the most promising current technology. RT-RPA is an isothermal amplification method in which the double stranded DNA denaturation and strand invasion that is typically achieved by heat cycling in PCR is instead accomplished by a cocktail of recombinase enzyme, single-stranded binding proteins, and ATP (*12*). RPA has potential advantages over other isothermal amplification technologies such as loop-mediated isothermal amplification (LAMP) (*13*) as it can be performed near ambient temperature (37-42°C). While several creative applications of LAMP technologies to detect COVID-19, the disease caused by SARS-CoV-2, have recently been developed and show promise (*14*–*18*), RT-RPA has been less explored.

## Reverse transcriptase choice can greatly affect recombinase polymerase amplification efficiency

We designed RPA primers to both the SARS-CoV-2 N gene and S gene (**fig. S1A and Table S1**) and quantified the amplification performance of a RT-RPA assay with ProtoScript II reverse transcriptase by qPCR (**fig. S1B**). The detection limit of this standard assay was poor, requiring between 100 and 300 RNA molecules for reliable detection (**Fig. 1A, fig. S1C** bottom panel). Some studies have used longer reaction times to partially counteract the poor yield of RT-RPA (*19*), but we set out to determine whether alternative approaches were possible.

**Figure 1.**
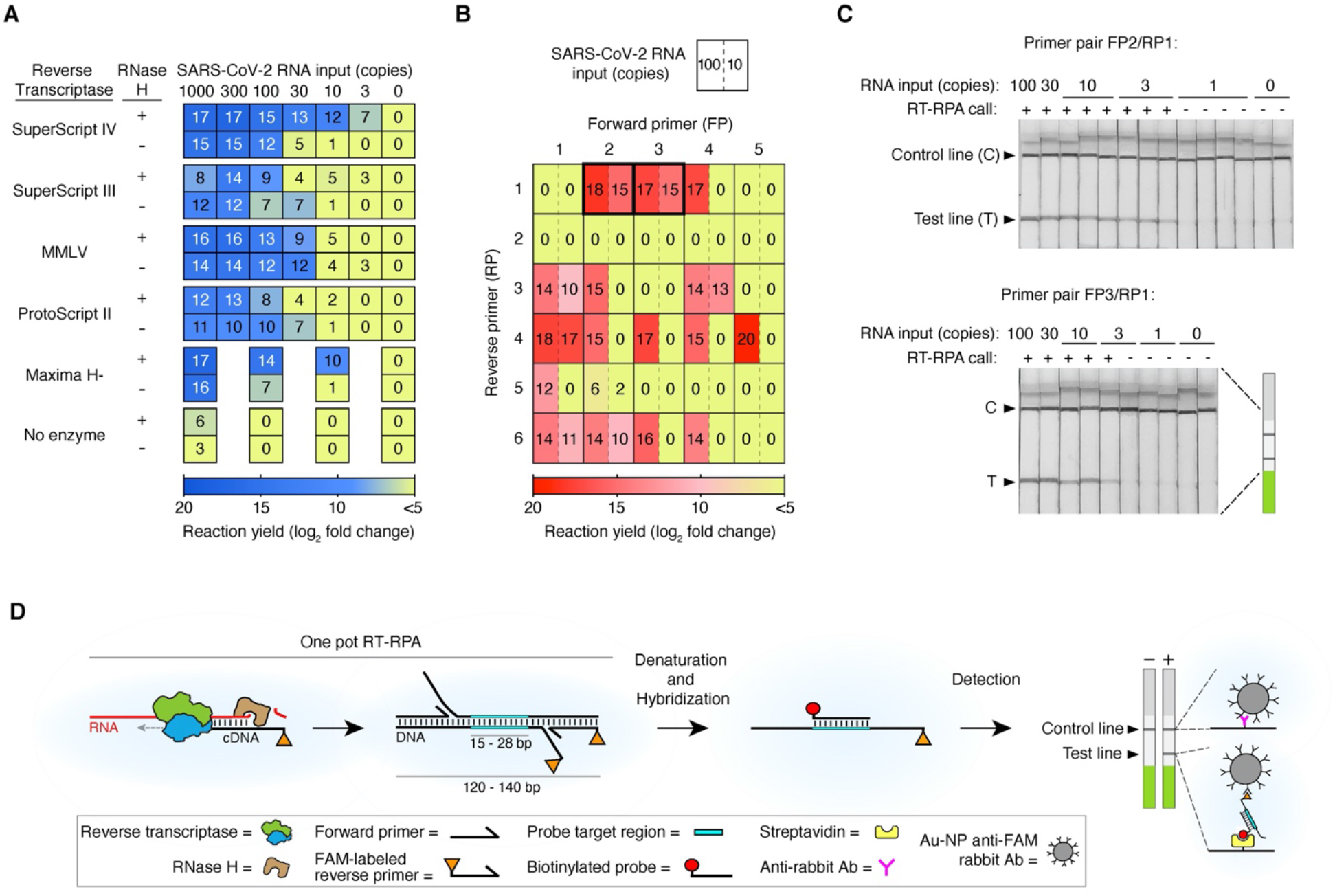
Development of FIND: an enhanced RT-RPA based assay for detection of SARS-CoV-2. **(A)** Screen for reverse transcriptase (RT) enzyme and effect of RNase H. SARS-CoV-2 RNA was amplified by RT-recombinase polymerase amplification (RT-RPA) using five different RTs with or without RNase H addition and the yield of each reaction was determined by quantitative PCR (qPCR). At least two biological and two technical replicates were used for each data point; numbers in each square represent mean log_2_ fold amplification. Samples labeled as zero yielded only non-specific amplification products. **(B)** Primer optimization screen. SARS-CoV-2 RNA was amplified by RT-RPA using forward and reverse primers specific to the S gene. The yield of each reaction was determined by qPCR using the same primer pair as for the RT-RPA reaction. Data represent mean log_2_ fold amplification from 2 technical replicates for each RNA input. **(C)** Lateral flow strip readout of RT-RPA reactions of SARS-CoV-2 RNA using primer pairs FP2/FAM-labeled RP1 and FP3/FAM-labeled RP1. All lateral flow strips contain a control (C) and test (T) band. **(D)** Schematic of FIND. Viral RNA is first copied to cDNA by RT, then degraded by RNase H. The cDNA product is amplified by RPA using a forward and a FAM-labeled reverse pair of primers specific to the target sequence. The amplified material is then denatured and hybridized to a biotinylated probe. Dual FAM- and biotin-labeled products are detected on lateral flow strips.

We reasoned that the poor performance of RT-RPA could either be due to a specific inhibitor of the RPA reaction from the RT (reverse transcription) reaction or to non-specific primer oligomerization products that could dominate the amplification reaction before the RT reaction occurs (**Data S1**). These possibilities are not mutually exclusive. As the RPA reaction is both fast and sensitive when DNA is used as an input (*12*, *20*), we further hypothesized that the product of the RT reaction, i.e. the RNA:DNA hybrid duplex, might inhibit the RPA reaction. We explored methods to circumvent both of these possible problems. To address the problem of kinetic interference by non-specific oligomerization, we screened multiple reverse transcriptases; and to attempt to remove interference from RNA:DNA hybrids, we introduced RNase H, which selectively degrades the RNA strand in these hybrids. Our tests showed that both RT enzyme choice and RNase H addition affected the sensitivity of the RT-RPA reaction, suggesting that both of our hypothesized mechanisms affect RT-RPA efficiency (**Fig. 1A, figs. S1C and D**). The best combination we identified was SuperScript IV reverse transcriptase with RNase H. The magnitude of the effect of the addition of RNase H was correlated with the intrinsic RNase H activity of the RT enzyme. Both SuperScript IV and Maxima H Minus reverse transcriptases are engineered to have minimal RNase H activity in order to improve their processivity, robustness, and synthesis rate (*21*), and we saw the largest effect of RNase H addition in RT-RPA reactions using these enzymes.

## Reducing non-specific primer reactions increases RT-RPA yield

In addition to the performance issues addressed above, non-specific amplification reactions of primer dimers can greatly inhibit the ability of RPA to amplify the sequence of interest (*10*). To determine whether primer choice affects the importance of these non-specific reactions, we designed forward and reverse primers to both the SARS-CoV-2 N gene and S gene (**fig. S1A**). Our primer designs avoided regions with strong homology to other coronaviruses including MERS and SARS-CoV, as well as HCoV-229E, HCoV-HKU1, HCoV-NL63, HCoV-OC43, which cause respiratory illnesses such as the common cold. We also avoided regions that have high variability across sequenced SARS-CoV-2 strains (**Data S2**). We screened all pairwise combinations of primers to find pairs that gave a high yield of the desired target sequence while minimizing the amount of non-specific amplicons. Primer pairs were screened by performing qPCR on diluted RT-RPA products so that both specific and non-specific reaction yield could be determined, using a modification of a method we previously developed (**Fig. 1B**) (*20*). Many pairs gave high levels of amplification at 100 molecules of input RNA, but only a small fraction of those yielded significant amplification products at 10 molecules of input RNA.

## An optimized RT-RPA reaction allows for simple detection

Our optimized RT-RPA assay’s product can be hybridized and detected with a commercial lateral flow assay (LFA) without further amplification. LFAs allow accurate read-out by eye by minimally trained personnel, and even opens up the possibility of home-based testing (*22*). We chose to use Milenia Biotec HybriDetect lateral flow test strips that contain a streptavidin band, an anti-Ig band, and carry gold nanoparticle-labeled anti-FAM antibodies for visualization. Based on the results shown in **Fig. 1B**, we selected two primer pairs that amplify part of the S gene, added a FAM label to the reverse primer, and hybridized the product amplicon to a biotinylated capture probe. Consistent with expectations from qPCR, both primer pairs reproducibly yielded bands with 10 input molecules, and one gave consistent bands with 3 input molecules (**Fig. 1C**). We called this optimized protocol FIND (Fast Isothermal Nucleic acid Detection) (**Fig. 1D**). Compared to the original RT-RPA assay using ProtoScript II RT, the detection limit for FIND was improved by several orders of magnitude (**fig. S1C**).

## FIND: a sensitive, specific, rapid test for SARS-CoV-2

FIND is highly sensitive and specific for SARS-CoV-2 N and S genes (**Fig. 2 and fig. S2**). The sensitive and specific assays were conducted by two independent groups, each of whom randomized the RNA input in a 96-well plate in a checkerboard pattern, then handed the blinded plate to the other group for testing by FIND (**fig. S2A**). For each gene, 52 positive samples were included with a concentration ranging from 100 molecules to 1 molecule of total RNA input (**Figs. 2A and C and figs. S2B and C**). The titer of the RNA dilutions was confirmed by RT-qPCR (**Fig. 2B and figs. S2D-F**). Strips were scored at ~20 minutes as this decreases the variability in band intensity that can be observed at low molecule input (**Fig. 2C and fig. S2B**). At or above 10 molecules of RNA input, 87 of 88 N gene samples and 88 of 88 S gene samples were accurately identified as SARS-CoV-2 positive (**fig. S2C**). Significant detection was achieved even as low as 3 (13 of 24 tests) or 1 (5 of 16 tests) molecules of RNA input. Critically, our assay is also highly specific, showing no cross-reactivity (0 of 80 tests) with 10,000 copies of RNA from other coronaviruses, i.e. MERS, SARS-CoV, CoV-HKU1, or CoV-229E. It also showed no crossreactivity with the 2009 H1N1 Influenza virus, a respiratory virus with similar initial clinical presentation (**Figs. 2A and C, fig. S2B**). For SARS-CoV and MERS, which have the highest target sequence identity with SARS-CoV-2 (91% and 66% respectively), cross-reactivity is dependent on probe choice; we observed cross-reactivity with MERS and SARS-CoV when a longer biotin-probe was used for detection (**fig. S2G**).

**Figure 2.**
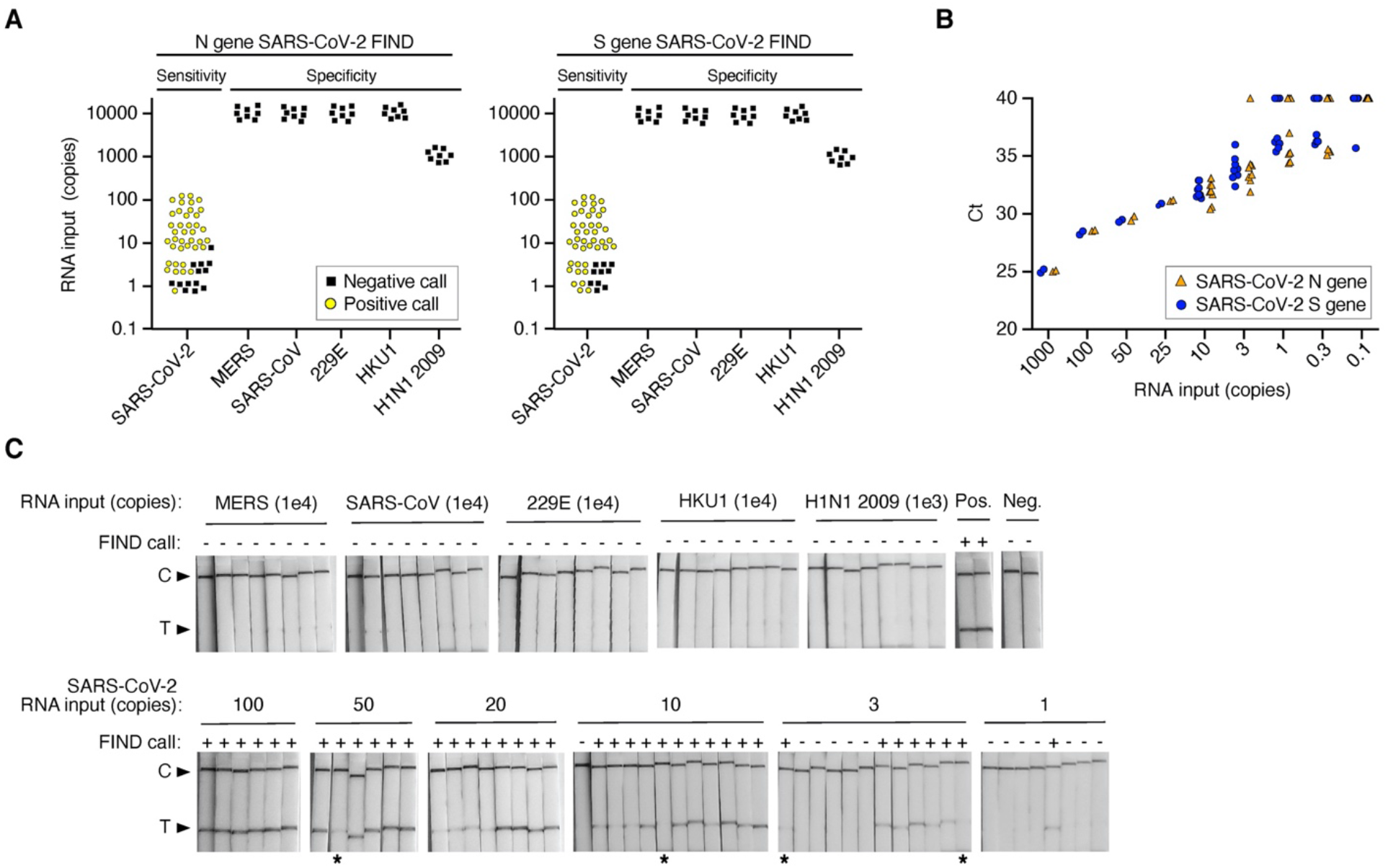
Sensitivity and specificity of RNA detection. **(A)** Summary of FIND test results for detection of RNA from SARS-CoV-2 or from other viruses. Synthetic full genome SARS-CoV-2 RNA was amplified by FIND using primers targeting the N or S gene and reactions were read out by lateral flow strip. The specificity of FIND was tested against either in vitro transcribed (IVT) RNA of the related viruses MERS and SARS-CoV, or IVT RNA of the common cold coronaviruses HCoV-HKU1 and HCoV-229E, or viral genomic RNA extracted from 2009 H1N1 Influenza. Data points represent positive (yellow) or negative (black) FIND tests for each sample tested and are staggered on both axes for visualization. **(B)** Quantification of the synthetic full genome SARS-CoV-2 RNA used as input in the FIND assay by RT-qPCR. Data are Ct values determined using a one-step commercial RT-qPCR assay using primers targeting either the N or S gene of SARS-CoV-2. Data points at Ct=40 represent non-specific or no amplification. N gene and S gene data are offset on the x-axis for visualization purposes. **(C)** Lateral flow strip readouts for all N gene data shown in (A). Individual strips are labeled with the test call made within 20 mins of detection (positive (+) or negative (-)). The positive (Pos.) FIND control is 1,000 copies of synthetic full genome SARS-CoV-2 RNA and the negative (Neg.) FIND control is a water-only input. Images taken for the purpose of display were allowed to dry which reduced the intensity of some weak bands (labeled with asterisks).

We developed an RNA extraction free lysis approach as RNA extraction from clinical samples has become a limiting factor as the global need for SARS-CoV-2 tests has increased. RNA extraction kits are currently hard to obtain, the process of extraction depends on skilled workers, and often involves equipment such as centrifuges. Heat-based lysis has shown promise as a way to rapidly lyse and inactivate viruses for use in diagnostic assays (*23*, *24*). To test whether heat-based sample lysis made viral RNA accessible for FIND, we initially used packaged reference viral particles, the AccuPlex SARS-CoV-2 verification panel (Seracare). We determined the relationship between temperature and viral lysis by heating for 5 minutes followed by RT-RPA then qPCR for quantification. The replication-deficient virus in the AccuPlex panel is lysed at ~75°C, a temperature that is likely similar to the temperature required to lyse wild-type SARS-CoV-2 (**Fig. 3A**) (*25*).

**Figure 3.**
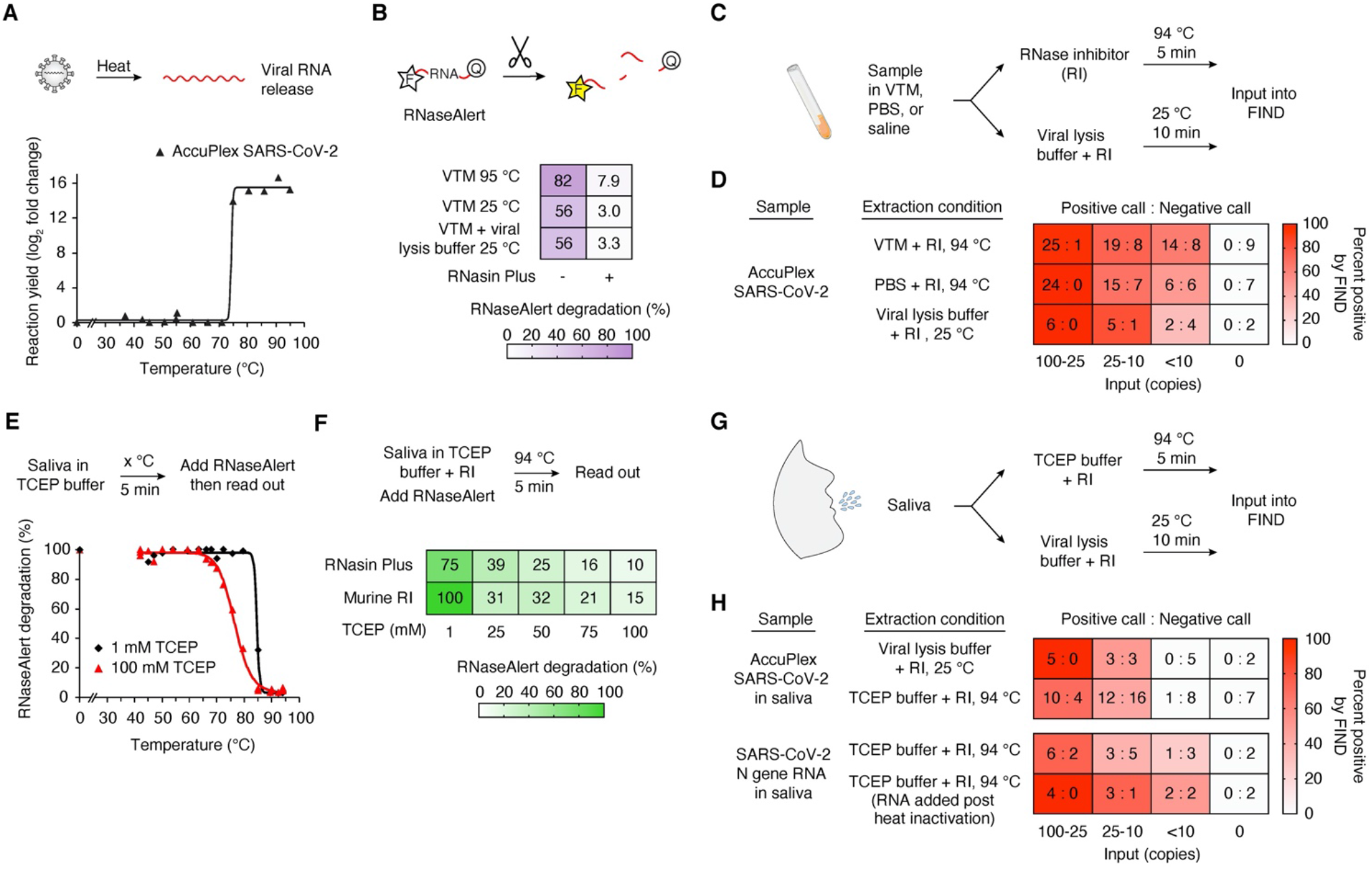
Lysis and detection of SARS-CoV-2 N gene from contrived samples. **(A)** Viral particle temperature lysis determination. AccuPlex packaged SARS-CoV-2 virus was diluted into TCEP buffer and heated for 5 min at the given temperature (see Methods). Released RNA was amplified by FIND and product formation was quantified by qPCR. **(B)** Detection of RNase activity of VTM. RNaseAlert was added to viral transport media (VTM) with or without the addition of RNasin Plus before heating for 5 min at 94°C or added to a 1:1 VTM and viral lysis buffer mix and incubating for 10 min at 25°C. Data represent the average of 4 technical replicates and were determined by normalizing the fluorescence intensity 10 mins after the heating step to a fully degraded control. **(C)** Schematic of sample processing of patient samples in VTM for input into FIND. **(D)** Heatmap displaying FIND test calls for detection of AccuPlex packaged SARS-CoV-2 lysed with conditions displayed in (C). AccuPlex packaged SARS-CoV-2 virus was mixed 1:1 with VTM, PBS, or viral lysis buffer and incubated as shown. All samples included RNasin Plus. Values represent the number of positive test calls: number of negative test calls for each condition. **(E**) Inactivation of RNase activity in saliva by TCEP and heat. Saliva was first mixed 1:1 with a buffer containing 1 mM (black) or 100 mM (red) TCEP and heated at the indicated temperature for 5 min. After cooling, RNaseAlert was added and degradation was assessed as in (B). **(F)** The combined activities of an RNase inhibitor and TCEP protect RNA from degradation in saliva. RNaseAlert was added to saliva diluted 1:1 with TCEP buffer containing an RNase inhibitor and treated as shown. RNAseAlert degradation was assessed an in (B). See additional data in **Fig. S3G**. **(G)** Schematic of sample processing of patient saliva samples for input into FIND. **(H)** Heatmap displaying FIND test calls for detection of SARS-CoV-2 RNA or AccuPlex packaged virus from saliva treated as displayed in (G). AccuPlex packaged SARS-CoV-2 virus or SARS-CoV-2 N gene IVT RNA were added to saliva and extracted as shown. Values represent the number of positive test calls: number of negative test calls for each condition.

RNase inhibitors prevent RNA degradation from nasopharyngeal swabs suspended in viral transport media (NP in VTM), the standard clinical sample. Our initial experiments using AccuPlex samples or in vitro transcribed RNA in VTM yielded poor signal intensities by FIND. Using RNaseAlert to measure RNase activity, we were surprised to find significant RNase activity in VTM (**Fig. 3B**). In an attempt to address this we tested TCEP, which has been used to inactivate RNases from saliva and urine (*26*). Unfortunately, TCEP and heat treatment of samples with VTM led to gelation, likely due to the presence of gelatin and bovine serum albumin in VTM (**fig. S3A**). As an alternative we tested RNasin Plus, a thermostable RNase inhibitor, which significantly protected RNaseAlert from degradation during heat-based lysis in VTM. For future compatibility with PON testing, we also tested a room temperature viral lysis buffer (Intact Genomic FastAmp^®^ Viral and Cell Solution for Covid-19 Testing) and found that RNaseAlert was protected from degradation in the presence of RNasin Plus (**Fig. 3B and figs. S3B and C**).

To confirm that this protocol is effective for patient samples, we tested heat-based lysis of NP-swabs in VTM in the absence and presence of RNasin Plus. We found the addition of RNasin Plus increased RNA yield by ~10-fold and significantly improved the sensitivity of FIND (**figs. S3B and C**). We also measured the sensitivity of FIND for AccuPlex viral particles diluted into VTM, PBS, or viral lysis buffer. Sensitivity in these simulated samples, which should more closely reflect what would be achieved from standard samples, was reduced by about 5-fold in comparison to RNA samples in water (**Figs. 2 and 3D**). Most patients during the initial active phase of infection deliver NP swabs with virus concentrations of >10^4^ per mL, well within our detection limit (*27*–*30*).

## Adaptation of FIND to detect virus in saliva

Given the bottleneck in NP swabs, there has been growing interest in testing saliva instead (*31*). Saliva is a challenging fluid due to the presence of mucins and RNases (*32*, *33*) which can degrade RNA and clog pipettes, leading to a high rate of failed experiments or false negatives. Nevertheless, the viral titer in saliva is sufficient for SARS-CoV-2 detection (*34*). To adapt FIND to saliva samples, we tested protocols that used TCEP, EDTA, and heat steps (*23*, *24*). The addition of the reducing agent TCEP was critical to decreasing the viscosity of saliva at all temperatures, but the inhibition of RNase activity by TCEP was not complete until the sample was heated above 85°C (**Fig. 3E**). Because SARS-CoV-2 viral particles lyse at around 75°C (*25*), the period when the sample is being heated from 75°C to 85°C offers a window in which released viral RNA might be degraded during sample preparation. Indeed, RNaseAlert is completely degraded even in the presence of 100 mM TCEP if it is added before the heat inactivation step, but protected if it is added after heat inactivation (**figs. S3D and E**). This distinction is critical as a common method of validating extraction-free saliva sample preparation protocols is to first heat-inactivate the sample and then add viral RNA to determine assay sensitivity (*15*). This method will overestimate assay sensitivity for saliva samples due to the inactivation of salivary RNases. Either murine RNase inhibitor or RNasin Plus helped protect RNA from degradation at low temperatures, with RNasin Plus being more effective at high temperatures (**fig. S3F**). The combination of RNAsin Plus and TCEP protects RNAseAlert from degradation during a heat lysis protocol (**Fig. 3F and fig. S3G**). Using this protocol (**Fig. 3G**) we detected SARS-CoV-2 signal in ~70% of samples with 25-100 AccuPlex viral particles in saliva (**Fig. 3H**), a reduction of 2 to 4-fold compared to the sensitivity of detection in VTM (**Fig. 3D**). We saw similar results with IVT SARS-CoV-2 RNA which represents the worst-case scenario for RNA degradation (**Fig. 3H**). Given that titers of SARS-CoV-2 in saliva are in the range of 10^4^ to 10^10^ copies per mL (*34*), this extraction protocol combined with FIND should be able to identify COVID-19 in a high proportion of infected patients, offering the potential for a high throughput, first pass screening approach that could be important in large-scale testing. We note that we have not yet tested FIND on actual saliva samples from infected individuals, as these are not readily available.

## Comparison of FIND with RT-qPCR tests on unextracted clinical samples

To demonstrate that FIND can detect SARS-CoV-2 in unextracted patient samples, we obtained 30 positive and 21 negative NP swabs from BocaBiolistics (**Data S3**). We processed the samples using our VTM heat lysis method (**Fig. 3C**) and used this unextracted input, in parallel, in FIND and in a one-step RT-qPCR assay (**Fig. 4A**). We validated our one-step RT-qPCR assay by benchmarking it against the standard CDC N1 RT-qPCR assay (**Fig. S4, see Methods**). All 21 negative samples were negative by FIND, in duplicate, confirming that the false positive rate for FIND is very low (**Figs. 4B-C, Data S3**). Of the 30 positive samples, 26 had signal by RT-qPCR; 4 may have suffered degradation during transit, see below. For each of the 26 samples that were positive by RT-qPCR, we estimated the number of copies of input RNA into FIND based on standard curves (**Data S3**). 20 samples had an input of at least 5 molecules of RNA; all 20 of these were positive by FIND in both repeats (**fig. S4A**). In 3 samples the input was between 1 and 4 copies; FIND was positive once, inconclusive once (one positive and one negative of two duplicates), and negative once. In 3 samples the input was less than 1 copy, and two of these three samples were inconclusive by RT-qPCR; FIND was negative twice and inconclusive once (**Fig. 4C**).

**Figure 4.**
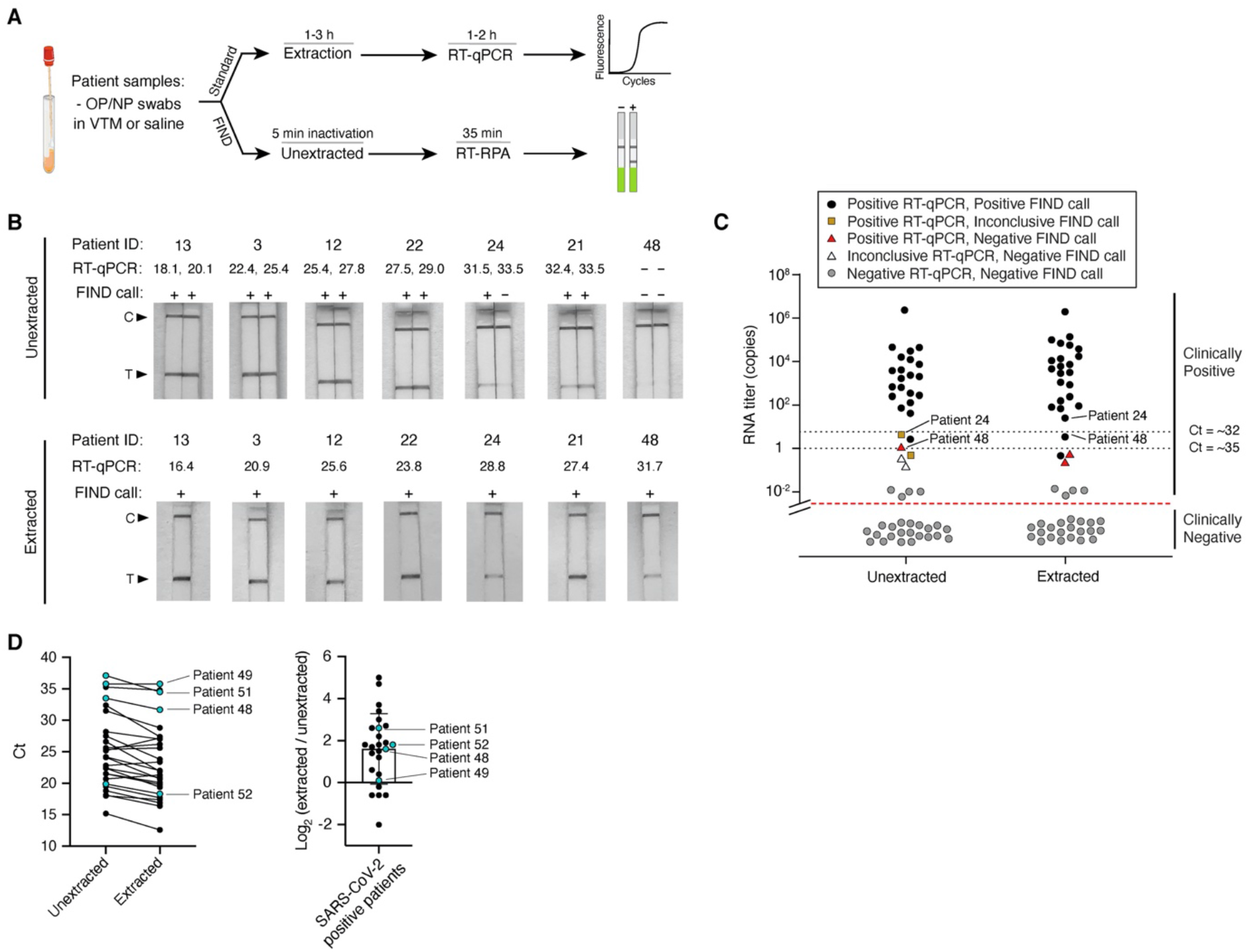
Detection of SARS-CoV-2 in clinical samples using FIND. **(A)** Schematic of the workflow for benchmarking FIND against RT-qPCR using patient samples. **(B)** Sampling of lateral flow strip readouts of SARS-CoV-2 N gene FIND tests of unextracted (Top) or extracted (Bottom) patient samples of known infection status. Unextracted patient samples were run in duplicates both by FIND (calls of positive (+) or negative (-) were made within 20 min of detection) and by one-step RT-qPCR (Ct values shown). See additional data in **Fig. S4A**. RNA was extracted from clinical samples according to standard procedure (see Methods) and was subsequently used as input to FIND and RT-qPCR. See additional data in **Fig. S4B**. **(C)** Summary of FIND test results of 51 patient samples and comparison to RT-qPCR. The y axis represents patient viral titer determined using a commercial one-step RT-qPCR assay from unextracted samples or extracted RNA samples with a standard curve. **(D)** (Left) Matched RT-qPCR Ct values of unextracted and extracted patient samples. (Right) Difference between extracted and unextracted Ct values for all patients. Patient samples were provided in multitrans media (black) or universal transport media (blue).

We repeated the FIND workflow on the S gene and obtained similar results to the N gene (**Fig. S5**). In all the patient samples, the RNA copy number of the S gene was on average 4-fold lower than that for the N gene (**fig. S5B**). This is puzzling, as the detected copy number for both genes are nearly identical from synthetic full genome SARS-CoV-2 RNA, AccuPlex viral particles, and IVT RNA controls (**figs. S5B-D**). One possible explanation for this result is that viral transcripts from cells contribute significantly to the total RNA detected. Because the N gene is expressed at up to 10-fold higher levels than the S gene in infected human cells (*35*), if NP swabs collect cells or cellular debris this would bias the observed N gene to S gene copy number ratio. This may be important for other assays as many COVID tests target ORF1ab, which is one of the lower expressed transcripts in human cells.

Sample degradation during storage could have substantially lowered the titer of some of our positive samples. Eight positive samples were stored in universal transport medium (UTM); only one of these had a titer above 2 RNA copies per μL. Four of these eight, although they were designated positive by BocaBiolistics, were negative by RT-qPCR and by FIND for both the N and S gene. To exclude the possibility that the low apparent titer was due to interference from UTM, we performed RNA extraction on all samples and then repeated RT-qPCR and FIND (**Fig. 4A**). Overall, RNA extraction increased RNA titer by ~5-fold, matching expectations given that 240 μL of initial sample was concentrated into 50 μL of final volume (**Figs. 4B-D**). Extraction of UTM samples did not differentially improve RT-qPCR or FIND results over VTM samples, excluding the possibility of interference from the medium and suggesting that the samples stored in UTM may have suffered RNA degradation. It is also possible that all 8 of these samples were unusually low titer on collection.

FIND gives concordant results with RT-qPCR in all extracted samples except those with extremely low titer. Of the 26 extracted samples that were detected as positive by RT-qPCR without extraction, 23 had at least 3 copies of input RNA, and all of these were positively identified by FIND (**fig. S4B**). Three samples had <1 copy of input RNA, of which one was identified by FIND. The four samples with undetectable signal by RT-qPCR before extraction were still negative by both RT-qPCR and FIND even with extraction. We note that modest changes in sample collection methods could make FIND even more sensitive. Currently, NP swabs are typically collected into 3 mL of VTM. Only a small fraction of this volume is used for detection assays. If instead swabs were resuspended in 150-200 μL of liquid, the volume required to cover the head of a swab, the input to FIND would become ~15-20-fold more concentrated without requiring an extraction protocol. This could make the sensitivity of FIND superior to current sample collection methods combined with RT-qPCR.

The FIND protocol reported here was developed and optimized in just under 3 weeks, with an additional 4 weeks for sample preparation, optimization, and patient sample acquisition. In future epidemics and pandemics, this process could be shortened to several days after standardizing sample preparation methods and primer design, and streamlining IRB and COMS approvals. A companion manuscript shows that the improvements in RT-RPA we developed in FIND also improve other detection approaches such as SHINE (*36*), allowing these assays to become 1-pot, closed-tube, fluorescent readout reactions. FIND addresses many of the problems of current SARS-CoV-2 testing methods: it is scalable, compatible with both swabs and saliva samples, can be performed in high throughput by minimally trained personnel in low-resource settings (**fig. S6**), and can be automated. FIND is capable of reliably detecting SARS-CoV-2 virus in patient samples that contain as low as 2 viral particles/μL, and is therefore fully adequate to detect infection during the period of peak transmission (*27*–*30*, *37*, *38*).

## Acknowledgments

We would like to thank Sam Keough, Raquel Arias-Camison, and Sandro Santagata for IRB and COMS help; the Sabeti lab, Dan Davidi, Cameron Myrhvold, Allon Klein, Sean Megason, Pam Silver, Galit Alter, Stephen Elledge, Brian Rabe, and Connie Cepko for thoughtful discussions; Hadley Weiss and Erika Olson from the Silver lab for N and S gene expression plasmids; Neal I. Lindeman and Michael K. Slevin from Brigham and Women’s Hospital for sample testing; and Laura Maliszewski, Peter Sorger, and Galit Lahav for structural support during the COVID mayhem. Funding: This work was supported by DARPA BRICS grant #HR001117S0029, the Quadrangle Fund for the Advancement and Seeding of Translational Research at Harvard Medical School (Q-FASTR), and the Massachusetts Consortium on Pathogen Readiness (MassCPR) and China Evergrande Group. JQ is supported by NSF GRFP. MS is supported by R01 GM120122-01. Author contributions: JQ, ZL, SAB, and MS conceived the study. JQ, ZL, SAB, CC, MEP, and BLG performed most key experiments and data analysis (supervised by MS). JMF and JZL were responsible for the patient collection and initial sample processing. RTI did the bioinformatic analysis. JQ, SAB, CC, ZL, MEP, BLG, RHW, and MS wrote the paper. All authors reviewed the manuscript. Competing interests: 3 patents have been filed U.S. Provisional Patent Application 63/993,521, 63/003,555, and 63/006,372. A company, Qtection LLC, has been launched to commercialize FIND and provide effective testing options to the public. MS is a founder of the company. Data and materials availability: All data are available in the article or the supplementary materials.

## Supplementary Materials

Materials and Methods

Figures S1-S6

Table S1

Data S1-S3

