## Supplemental materials for "An enhanced isothermal amplification assay for viral detection"

**This PDF file includes:**

Materials and Methods
Supplementary Text
Figures S1 to S6
Table S1

**Other Supplementary Materials for this manuscript include the following:**

Data S1 to S3

### Materials and Methods

RNA template preparation

SARS-CoV-2, SARS-CoV, and MERS N gene containing plasmids were obtained from IDT (2019-nCoV Plasmid Controls). HCoV-229E and HCoV-HKU1 N gene, SARS-CoV-2, SARS-CoV, MERS, HCoV-229E and HCoV-HKU1 S gene were synthesized by Twist Bioscience. All genes were cloned into a T7 promoter expression plasmid. To produce RNA template, in vitro transcription was performed with NxGen® T7 RNA Polymerase (Lucigen #F88904-1) according to the manufacturer's suggested protocol with minor modifications. Final concentrations of the reaction mixture components were 50 units T7 RNA polymerase, 1× reaction buffer, 625 µM NTPs, 10 mM DTT, 500 ng of linearized plasmid template, and RNase-free water to a final volume of 20 µL per reaction. After 10 h at 37 ˚C, 4 units of DNase I (NEB #M0303S) was added and reactions were further incubated for 10 min at 37 ˚C. DNase I was heat inactivated by adding EDTA (5 mM final) and heating at 75 ˚C for 10 min. RNA was purified by RNAClean XP (Beckman Coulter) at 0.6× the volume of the reaction, washed twice with 80% EtOH, then eluted into 20 µL RNase-free water. The size and quality of the RNA product was checked by Bioanalyzer (Agilent) after denaturation at 70 ˚C for 2 min to unfold any RNA structure; all samples were determined to contain the correct RNA product.

Reverse transcription and RNase H screen

FIND assay master mixes targeting SARS-CoV-2 N gene as described below with or without addition of RNase H (NEB) and without reverse transcriptase. The following RT enzymes were added to aliquots of the master mixes: SuperScript III (ThermoFisher), SuperScript IV (ThermoFisher), MMLV (Moloney Murine Leukemia Virus RT, NEB), ProtoScript II (NEB), Maxima H Minus RT (ThermoFisher). All enzymes were added at 20 U per reaction. N gene IVT RNA diluted with $H_2O$ was used as input to the reactions. Post isothermal amplification, samples were diluted 1:400 in water and products were detected by qPCR using primers JQ289 and JQ223. Specific products were distinguished from primer dimers by analyzing the melting temperature of the qPCR products. The average Ct value of all water control reactions representing primer dimer was used as a baseline to determine the reaction yield (yield = Ct (average water controls) – Ct (specific reaction)).

Primer Oligomerization Products

Four N gene forward primers (JQ217, CCMS041, CCMS047, and CCMS051) were paired with the reverse primer JQ224. These four primer pairs as well as JQ217 + JQ223 were used in FIND assays with a water-only sample input. FIND assays were incubated at 42 ˚C for 10 min. Amplification products were cleaned up using RNA Clean XP (Beckman Coulter) at 2.5× concentration and eluted in 20 µL of nuclease-free water. Purified products were cloned using the Zero Blunt TOPO PCR Cloning Kit (Thermo Fisher Scientific) according to the manufacturer’s instructions and Sanger sequenced. The identity of cloned products was determined by first aligning the sequences to the vector sequence using Samtools (v1.9) with an allowed multimapping of k=10000. The sam files were then visualized in IGV (v2.6.2) where the direction, sequence, and copy number of primer oligomers were manually annotated. In Table S2, the direction of the primer indicates a forward or reverse direction with respect to the vector, while the space sequence is the string of nucleotides between two primers. Overlap indicates the primers were overlapping with respect to the vector, "*" indicates there was no space between the primers, and listed nucleotides indicate the sequence between two primers.

SARS-CoV-2 FIND assay primer sequence alignment
To calculate the percent identity between the SARS-CoV-2 N and S gene primers and the analogous sites in other betacoronaviruses, the RefSeq entries for SARS-CoV-2, SARS-CoV, MERS, HCoV-229E, HCoV-NL63, HCoV-OC43, and HCoV-HKU1 were obtained from NCBI. The sequences were then compared using the EMBL-EBI web tool Clustal Omega to identify indels and mismatches. The subsequences for the forward and reverse primers for both the N gene and the S gene were then located within the SARS-CoV-2 sequence, and the number of mismatches with the antagonist betacoronavirus sequence was tallied. The percent identity was then calculated by dividing the number of matching bases by the length of the primer sequence.
To calculate the number of mismatches between FIND assay primer and probe sequences and known SARS-CoV-2 variants, the full set of all available SARS-CoV-2 genomes were downloaded from NCBI and were arranged into a single fasta file. This dataset was then converted into a BLAST database using the BLAST+ (v2.6.0) tool and then queried by each of the sequences for the N and S gene FIND assay. The output from BLAST was then coalesced and filtered to remove any incomplete or partial genomes using R (v.4.0).

Primer screening
Regions of low homology between SARS-CoV-2 and both SARS-CoV and MERS were identified by sequence alignment and were used as target sequences for the biotinylated probe. Unlabeled forward and reverse primers were designed to amplify a region of 100-200 nt encompassing the target sequence. Combinations of forward and reverse primers were screened by testing amplification at low RNA input. Reactions were prepared as described below and S gene IVT SARS-CoV-2 RNA was used as input. Post amplification, samples were diluted 1:625 in water and products were detected by qPCR using the same primer pair as for isothermal amplification. Specific products were distinguished from primer dimers by analyzing the melting temperature of the qPCR products. Reactions using water only as template were used to identify primer dimer melting temperatures. All reactions with 10 or 100 copies input but leading only to the formation of primer dimers were labeled as having a reaction yield of zero. The Ct of all reactions leading to specific product formation were converted into an estimated reaction yield by subtracting the raw Ct from the Ct of the lowest specific reaction (for the S gene screen the lowest specific Ct was at 25). Primer pairs with high reaction yields at both 100 and 10 copies input were tested in a secondary screen and the top two primer pairs were subsequently tested by FIND as described above.

FIND assay
Isothermal amplification reactions were based on the TwistAmp Basic RPA Kit (TwistDx) with added modifications described below. Each lyophilized pellet was resuspended in a solution of 38 µL rehydration buffer (TwistDx), 1 µL RNase H (5U/µL) (NEB), 0.5 µL SuperScript IV RT (200 U/µL) (ThermoFisher Scientific), and 0.5 µL of forward and reverse primer mix each at 50 µM (N gene, JQ217+JQ235; S gene, CCMS055+CCMS073). This mix was then activated by addition of 1 µL 700 mM magnesium acetate followed by thorough mixing with a pipette. Reactions were prepared by dispensing 8 µL of master mix and 2 µL of input template (RNA, Accuplex virus, or patient samples) per reaction well, mixing the reaction by pipetting, and incubating at 42 ˚C for 25

min. A hybridization mix was prepared by combining 1 vol biotinylated probe at 5 µM (N gene, JQ241 or JQ312; S gene, CCMS069) with 19 vol 10 mM Tris pH 8. 20 µL of hybridization mix was added to each reaction, and samples were heated at 94 ˚C for 3 min followed by a cooling step at room temperature for 3 min. 50 µL of Milenia GenLine Buffer (Milenia Biotec) was added to each reaction, mixed by pipetting, and a lateral flow strip (Milenia HybriDetect) was added. Lateral flow strip signals can be detected and imaged starting 3 min after addition of the strip to the hybridized reaction. Test results were called or imaged within 30 min of strip addition since background bands at the test line can appear over time and low signal test bands can lose intensity as the strip dries.

qPCR and RT-qPCR

SYBR green qPCR reactions were prepared in 10 µL reaction volume using PowerUp SYBR Green PCR Master mix (Thermo Fisher Scientific), 2 µL sample, and 0.4 µM of primers (JQ217 + JQ223 for N gene or CCMS055 + CCMS067 for S gene unless otherwise mentioned). RT-qPCR reactions were prepared in 10 µL reaction volume using the Luna Universal One-Step RT-qPCR kit (NEB), 2 µL sample, and 0.4 µM of primers following the manufacturer's instructions. The CDC one-step RT-qPCR assay used to benchmark our in house RT-qPCR was performed using the Luna Universal Probe One-step RT-qPCR kit (NEB) and N1 probe/primer mix against SARS-CoV-2 from IDT (2019-nCoV CDC EUA Kit) (**Fig. S4D**). Reactions were prepared according to the manufacturer's instructions following the CDC protocol. qPCR and RT-qPCR reactions were monitored on either a Bio-Rad C1000 Touch Thermo Cycler (Bio-Rad) or QuantStudio 6 Real Time PCR system (Thermo Fisher Scientific).

Sensitivity and specificity of FIND with RNA input

Data presented in Figure 2 was generated as a blinded and randomized experiment. Synthetic full genome SARS-CoV-2 RNA (Twist Bioscience) was used as RNA template for FIND assay on SARS-CoV-2. For the cross-reactivity samples, a single dilution series of RNA input was prepared by mixing at equimolar ratio N and S gene IVT RNA products for each of: SARS-CoV, MERS, HCoV-HKU1, and HCoV-229E. Genomic 2009 H1N1 Influenza (ATCC) was also serially diluted for input to the assay. All dilutions series were made in water and were adjusted for a 2 µL input into the FIND assay. Two independent groups prepared fully randomized 96-well PCR plates in a checkerboard pattern using those dilutions (**Fig. S2A**). Each group then used the other group's randomized plate as input to FIND tests targeting either the N gene or the S gene of SARS-CoV-2 performed as described above. All RNA stocks used in these tests were validated by testing dilution series in a one-step RT-qPCR as described above (**Fig. 2B and Figs. S2D-F**).

RNaseAlert tests with viral transport media (VTM) and saliva

The RNaseAlert substrate (IDT) was used at 2 µM to assess the RNase activity of saliva and VTM (BD, universal viral transport medium #2220220). Fluorescence intensity was determined using an excitation of 485 nm and emission of 528 nm over the course of 10-60 min in a 96-well plate reader (Synergy H1 Plate Reader, BioTek). In general, the degradation of the RNaseAlert substrate was assessed after 10 minutes and fluorescence intensities were averaged over 3 time points and reported normalized to a fully degraded control.

RNasin Plus (Promega) was added to VTM to a final concentration of 1 U/µL and was incubated for 5 min at 25 ˚C before addition of RNaseAlert. When needed, viral lysis buffer (FastAmp Viral and Cell solution, Intact Genomics) was added 1:1 (v/v) to VTM. TCEP buffer (20 mM Tris pH

8, 10 mM EDTA pH 8, TCEP 1-100 mM) was prepared as a 2× solution and was mixed 1:1 with saliva. RNase inhibitor (RI) was added to 1 U/µL final concentration as shown. For spike-in controls, RNase A (Lucigen) was added to 0.25 µg/µL final concentration. Saliva obtained from two healthy donors was pooled and adjusted to 1 mM TCEP to reduce viscosity. Aliquots of a single pooled sample stored at -20 ˚C were used for all assays.

Virus extraction

The AccuPlex SARS-CoV-2 verification panel (Seracare) containing the N gene, E gene, ORF1a, and RdRp was used as a surrogate to SARS-CoV-2 to optimize the full processing of clinical samples. To determine the temperature lysis of AccuPlex SARS-CoV-2, virus at 1e5 copies/mL was diluted 1:1 in 2× lysis buffer (final: 10 mM Tris HCl pH 8, 5 mM EDTA pH 8, 100 mM TCEP, 1 U/µL RNasin Plus), then incubated for 5 min at a temperature between 55 ˚C and 95 ˚C in 5 ˚C increments. 2 µL of each condition was used as input into FIND reactions targeting SARS-CoV-2 N gene as described above. Post amplification, samples were diluted 1:200 in water and products were detected by qPCR using primers JQ289 and JQ223. Specific product formation was distinguished from primer dimer formation by analyzing the melting temperature of the qPCR products and comparison to a water control. The average Ct value of all water control reactions representing primer dimer was used as a baseline to determine the reaction yield (yield = Ct (average water controls) – Ct (specific reaction)).

Detection of AccuPlex SARS-CoV-2 in contrived samples

AccuPlex SARS-CoV-2 was extracted using conditions mimicking patient sample processing. FIND assays targeting SARS-CoV-2 N gene were performed as above. For extraction in VTM and PBS, AccuPlex SARS-CoV-2 at 100 copies/µL was serially diluted 1:1 (v/v) in either VTM or PBS containing a final concentration of 1 U/µL RNasin Plus. After heating at 94 ˚C for 5 min, samples were kept on ice before being used as input into FIND. For extraction in viral lysis buffer at 25 ˚C, AccuPlex SARS-CoV-2 at 100 copies/µL was serially diluted 1:1 (v/v) in viral lysis buffer (FastAmp Viral and Cell solution (Intact Genomics)) adjusted with RNasin Plus to 1 U/µL. After 10 min at 25 ˚C, samples were kept on ice before being used as input into FIND. For extraction of virus in samples containing saliva, 2 vol of AccuPlex SARS-CoV-2 virus at 100 copies/µL was mixed with 1 vol of pooled saliva and 1 vol of 4× TCEP buffer + RI. TCEP buffer + RI was prepared such that final buffer concentrations in the sample were 10 mM Tris HCl pH 8, 5 mM EDTA pH 8, 100 mM TCEP and 1 U/µL RNasin Plus. Lower input samples were prepared by serial dilution with 1:1 (v/v) saliva in 2× TCEP buffer. After heating at 94 ˚C for 5 min, 1/10 vol of 1M $H_2O_2$ was added and samples were incubated at 25 ˚C for 10 min. Saliva samples were diluted 1:1 with water and kept on ice before being used as input into FIND. For extraction of virus from saliva with viral lysis buffer, 1 vol of AccuPlex SARS-CoV-2 virus at 100 copies/µL was mixed with 1 vol of pooled saliva and 2 vol of viral lysis buffer adjusted to 2 U/µL RNasin Plus. Lower input samples were prepared by serial dilution with 1:3 (v/v) viral lysis buffer + RI mixed with saliva. For samples with SARS-CoV-2 RNA, saliva was mixed 1:1 with 2× TCEP buffer + RI. After 5 mins at 25 ˚C, N gene IVT SARS-CoV-2 RNA was spiked into saliva in TCEP buffer and lower input samples were prepared by serial dilution on ice. After heating at 94 ˚C for 5 min, 1/10 vol of 1 M $H_2O_2$ was added and samples were incubated at 25 ˚C for 10 min. Samples were diluted 1:1 with water and kept on ice before being used as input into FIND. For RNA added post heat inactivation, a similar protocol was followed using saliva mixed 1:1 with 2× TCEP buffer + RI that was pre-incubated for 5 min at 94 ˚C.

Clinical samples

A cohort of nasal swab patient samples was purchased from BocaBiolistics, FL containing 30 SARS-CoV-2 positive samples and 21 SARS-CoV-2 negative samples. Samples were thawed on ice and 40 µL aliquots were made and subsequently stored at -80 ˚C. At the time of testing, sample aliquots were thawed and RNasin Plus was added to a final concentration of 1 U/µL. The samples were placed on a heat block set to 99 ˚C for 5 min for virus inactivation and lysis. After cooling, samples were spun down and transferred to a 96-well DNA LoBind plate (Eppendorf). 2 µL of the inactivated sample was used as input into FIND or into RT-qPCR reactions targeting both the N and S gene of SARS-CoV-2 (Supplementary Table 4). GAPDH was used as a control in RT-qPCR reactions. All patient sample tests included a positive control consisting in 100 copies of synthetic full genome SARS-CoV-2 RNA (Twist Bioscience) and a water only negative control.

Standard RNA extraction from clinical samples

Virions were pelleted by centrifugation at approximately 21,000 ×g for 2 h at 4 ˚C. The supernatant was removed and 750 µL of TRIzol-LS™ Reagent (ThermoFisher) was added to the pellets and then incubated on ice for 10 min. Following incubation, 200 µL of chloroform (MilliporeSigma) was added, vortexed, and incubated on ice for 2 min. Phases were separated by centrifugation at 21,000 ×g for 15 min at 4 ˚C, and subsequently the aqueous layer was removed and treated with 1 vol isopropanol (Sigma). GlycoBlue™ Coprecipitant (15 mg/mL) (ThermoFisher) and 100 µL 3M Sodium Acetate (Life Technology) were added to each sample and incubated on dry ice until frozen. RNA was pelleted by centrifugation at 21,000 ×g for 45 min at 4 ˚C. The supernatant was discarded and the RNA pellet was washed with cold 70% ethanol. RNA was eluted in 50 µL of DEPC-treated water (ThermoFisher).

Quantitative SARS-CoV-2 RT-qPCR Assay

Levels of SARS-CoV-2 RNA in extracted samples were detected using the US CDC 2019-nCoV_N1 primers and probe set. Each reaction contained extracted RNA, 1× TaqPath™ 1-Step RT-qPCR Master Mix, CG (ThermoFisher), 500 nM of each the forward and reverse primers, and 125 nM of probe. Viral copy numbers were quantified using N1 qPCR standards to generate a standard curve. The assay was run in triplicate for each sample and two non-template control (NTC) wells were included to confirm there was no contamination. Quantification of the Importin-8 (IPO8) housekeeping gene RNA level was performed to determine the quality of sample collection. An internal virion control (RCAS) was spiked into each sample and quantified to determine the efficiency of RNA extraction and qPCR amplification.

In house RT-qPCR data was converted from Ct values to copies/mL by direct comparison to the CDC RT-qPCR quantitation. In short, the Ct values from the in house RT-qPCR were plotted against the CDC RT-qPCR Ct values which yielded a linear relationship, R2>0.99, with a slope within error of 1, confirming that the amplification dynamics of both primer sets were similar. The relationship was then re-fit with the slope set to 1 which yielded a line, R2>0.99, with an intercept of between 33 and 34 (95% confidence interval). This fit was then used to directly convert Ct from the in house qPCR to viral copies/µL.

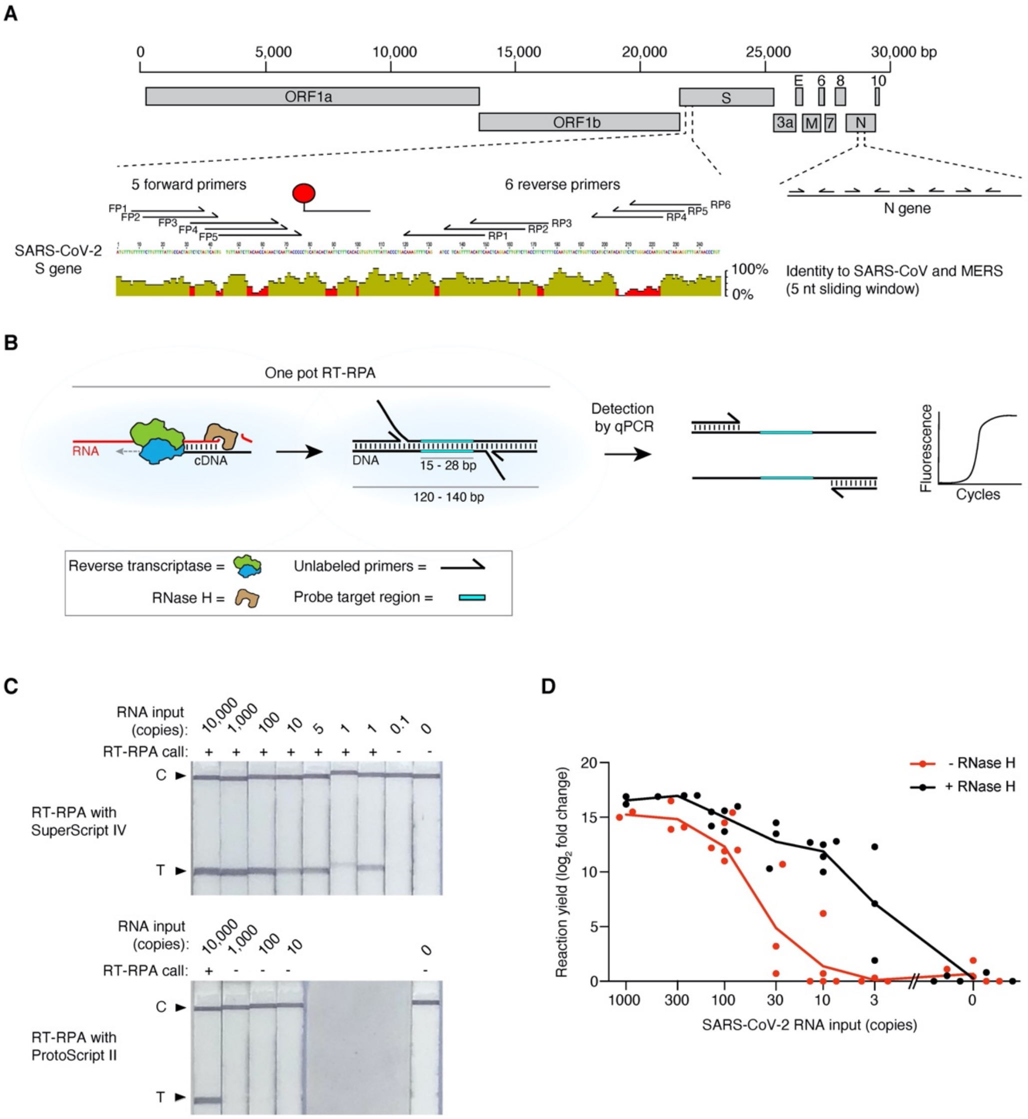


**Figure S1. Development of FIND. (A)** Organization of the SARS-CoV-2 genome and location of regions in the S and N genes targeted by FIND. Detailed mapping of the binding site of all forward and reverse primers tested in the primer optimization screen and of the biotin hybridization probe was shown for S gene only for display purposes. SARS-CoV-2 was aligned to the closely related SARS-CoV and MERS to identify regions of low homology which were targeted by primers and hybridization probes used in the assay. **(B)** Schematic of the workflow used for optimization of FIND. The cDNA product amplified by recombinase polymerase amplification (RPA) using forward and reverse unlabeled primers was quantified in a subsequent qPCR assay. **(C)** Comparison of the performance of SuperScript IV and ProtoScript II. In vitro transcribed (IVT) N gene SARS-CoV-2 RNA was amplified by RT-RPA and reactions were read out on a lateral flow strip. **(D)** IVT N gene SARS-CoV-2 RNA was amplified by RT-RPA with or without RNase H addition and the yield of each reaction was determined by quantitative PCR. Data represent the average yield of two technical replicates and is staggered on the x axis for visualization purposes.

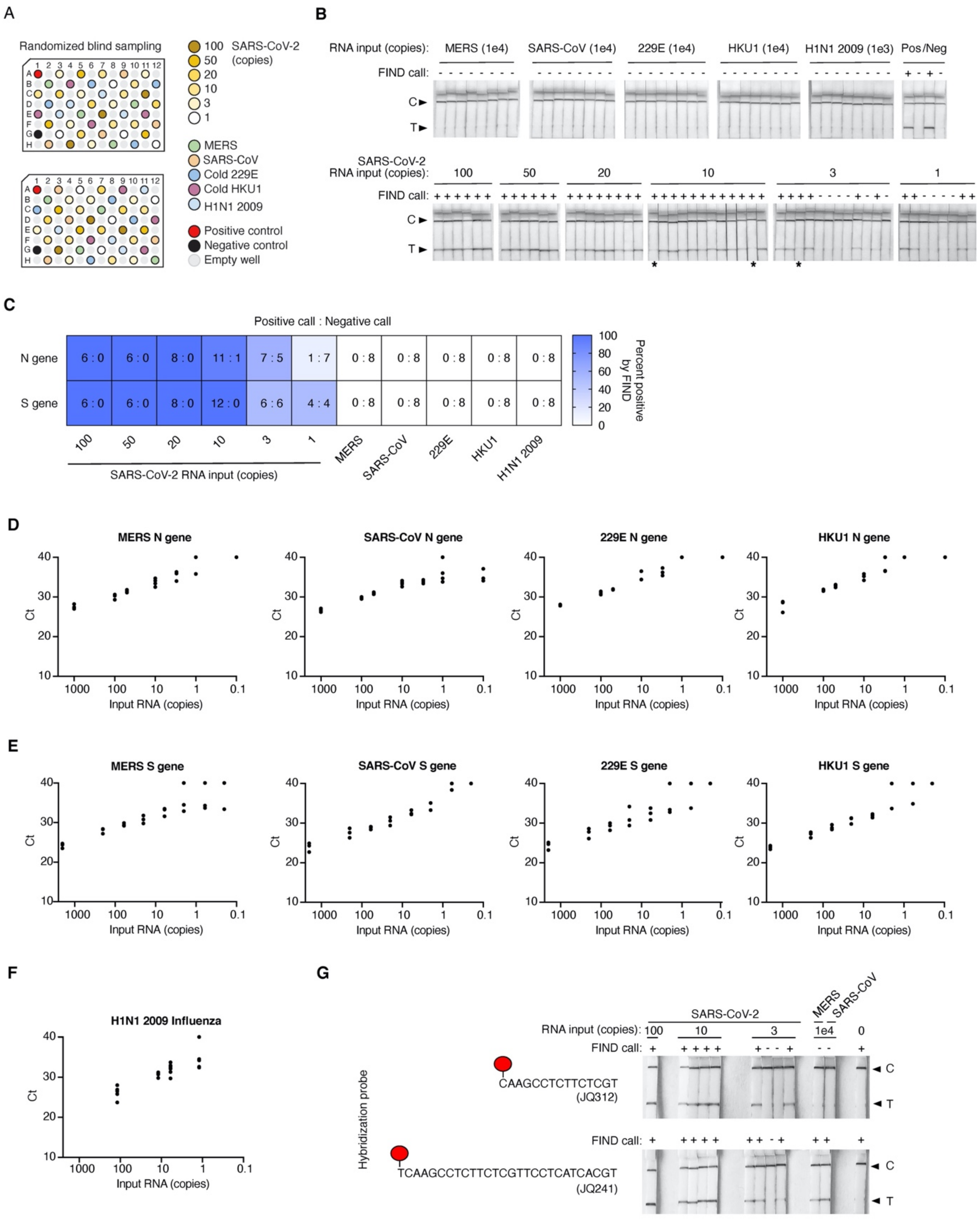


**Figure S2. Sensitivity and specificity of RNA detection. (A)** Blinded and randomized plate layout used in FIND assays used for generation of the data displayed in **Fig. 2A**. **(B)** Lateral flow strip readouts for the S gene dataset displayed in **Fig. 2A**. Individual strips are labeled with the test call made within 20 mins of detection (positive (+) or negative (-)). The positive (Pos.) FIND control is 1,000 copies of synthetic full genome SARS-CoV-2 RNA and the negative (Neg.) FIND control is a water-only input. Images taken for the purpose of display were allowed to dry which reduced the intensity of some weak bands (labeled with asterisks). **(C)** Heatmap displaying the rate of FIND test calls for detection of RNA from SARS-CoV-2 or from other viruses as shown in **Fig. 2A**. Values represent the number of positive test calls : number of negative test calls for each condition. **(D-E)** RT-qPCR quantification of in vitro transcribed (IVT) RNA from MERS, SARS-CoV, HCoV-229E, and HCoV-HKU1 used as specificity control tests in FIND; N gene shown in (D) and S gene in (E). **(F)** RT-qPCR quantification of RNA extracted from 2009 H1N1 Influenza. **(G)** Comparison of the specificity and sensitivity of two hybridization probes targeting SARS-CoV-2 N gene. IVT RNA from SARS-CoV-2, MERS, or SARS-CoV was amplified by FIND. After splitting the reactions in half and hybridizing with a biotinylated probe as shown, each reaction was read out on a lateral flow strip. Individual strips are labeled with the test call made within 20 mins of detection (positive (+) or negative (-)).

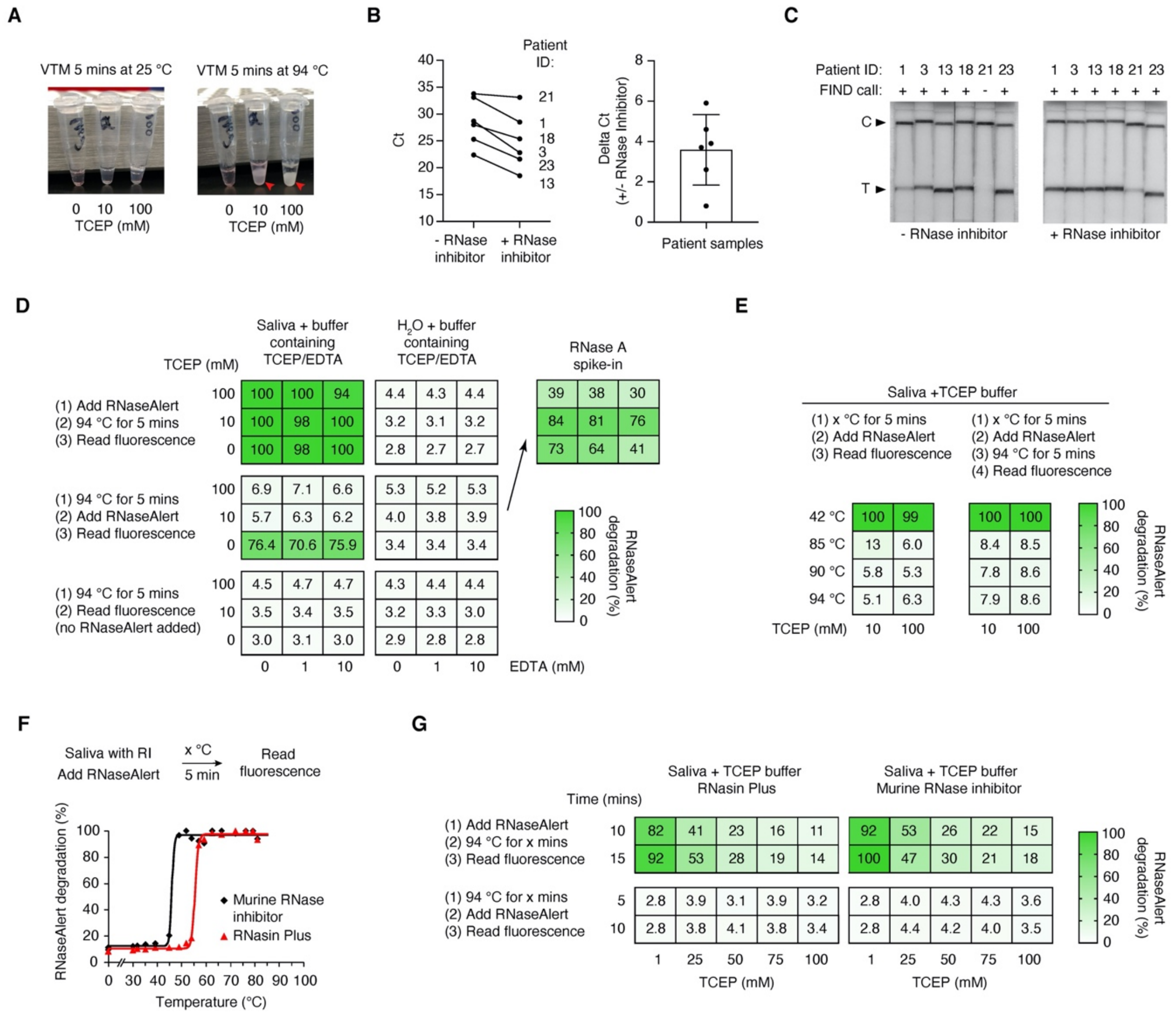


**Figure S3. Optimization of sample processing conditions for detection of SARS-CoV-2 in clinical samples. (A)** Heating VTM in presence of TCEP leads to formation of a gelatinous substance (highlighted by red arrowhead). **(B)** Addition of RNase inhibitor to patient samples prior to heat inactivation increases the RNA titer as quantified by RT-qPCR. Unextracted known positive patient samples were heat inactivated for 5 min at 94°C with or without RNasin Plus. Viral RNA was quantified using a commercial one-step RT-qPCR assay. (Left) Ct values for matched samples with and without RNase inhibitor. (Right) Difference between Ct values in all matched samples. Error bars represent +/- 1 standard deviation. **(C)** Addition of RNase inhibitor to patient samples prior to heat inactivation increases the signal of the FIND assay. Heat inactivated samples prepared in (B) were tested using FIND. **(D)** TCEP and heat (not EDTA) are required to inactivate the RNase activity in saliva as determined using RNaseAlert assays. Saliva (or water control) was mixed 1:1 with a buffer containing TCEP and EDTA as shown. RNaseAlert was added and the sample was heated as indicated. RNase A was added to a set of water samples post addition of RNaseAlert as control. Data represent the average of 2 technical replicates and was determined from the fluorescence signal 10 mins after the heating step normalized to a fully degraded control. **(E)** TCEP and heat irreversibly inactivate the RNase activity of saliva. Saliva was mixed 1:1 with a buffer containing TCEP and was processed as indicated. Data represent the average of 3 technical replicates and was determined as in (D). **(F)** RNase inhibitors protect RNA against degradation in saliva at low temperature only. Saliva was mixed 1:1 with a buffer containing an RNase inhibitor as shown. RNaseAlert was added and the sample heated as indicated. Data represent the fluorescence intensity 10 mins after the heating step normalized to a fully degraded control. **(G)** The combined activities of an RNase inhibitor and TCEP protect RNA from degradation in saliva (Additional data for **Fig. 3F**).

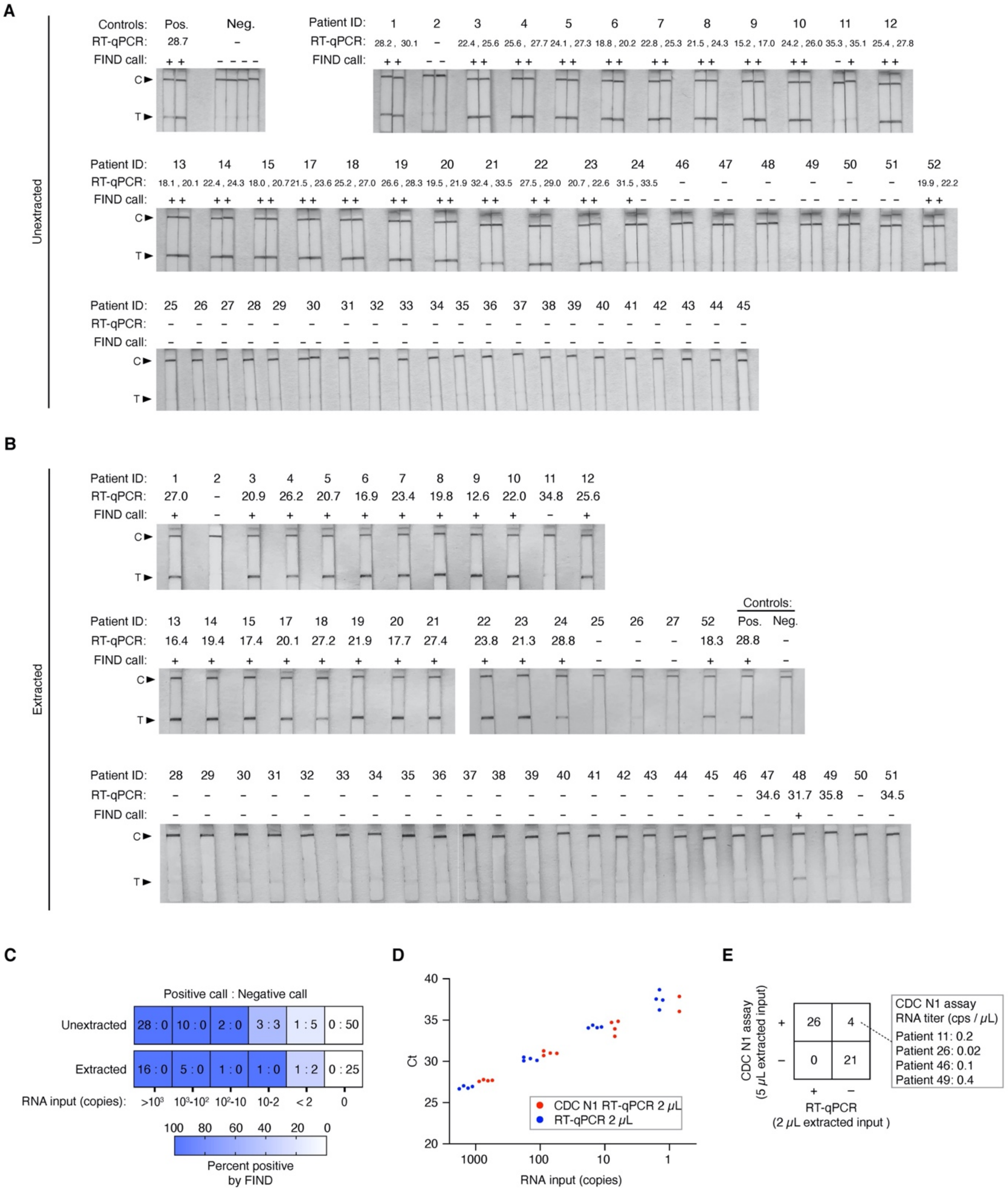


**Figure S4. Detection of SARS-CoV-2 from unextracted and extracted clinical samples. (A-B)** Lateral flow strip readouts of all FIND tests from unextracted (A) and extracted (B) patient samples summarized in **Fig. 4D**. Individual strips are labeled with the FIND test call made within 20 mins of detection (positive (+) or negative (-)). The positive (Pos.) FIND control is 100 copies of synthetic full genome SARS-CoV-2 RNA and the negative (Neg.) FIND control is a water-only input. **(C)** Heatmap displaying the rate of positive FIND test calls for detection of SARS-CoV-2 N gene from the 51 patient samples as shown in **Fig. 4D** binned by RNA input determined by one-step RT-qPCR. Values represent the number of positive test calls : number of negative test calls for each condition. **(D)** Our one-step RT-qPCR assay was validated against the CDC N1 RT-qPCR assay using synthetic SARS-CoV-2 RNA as input. **(E)** Comparison between the sensitivity of the CDC N1 RT-qPCR assay ran on 5 μL extracted sample and our one-step RT-qPCR assay ran on 2 μL extracted sample.

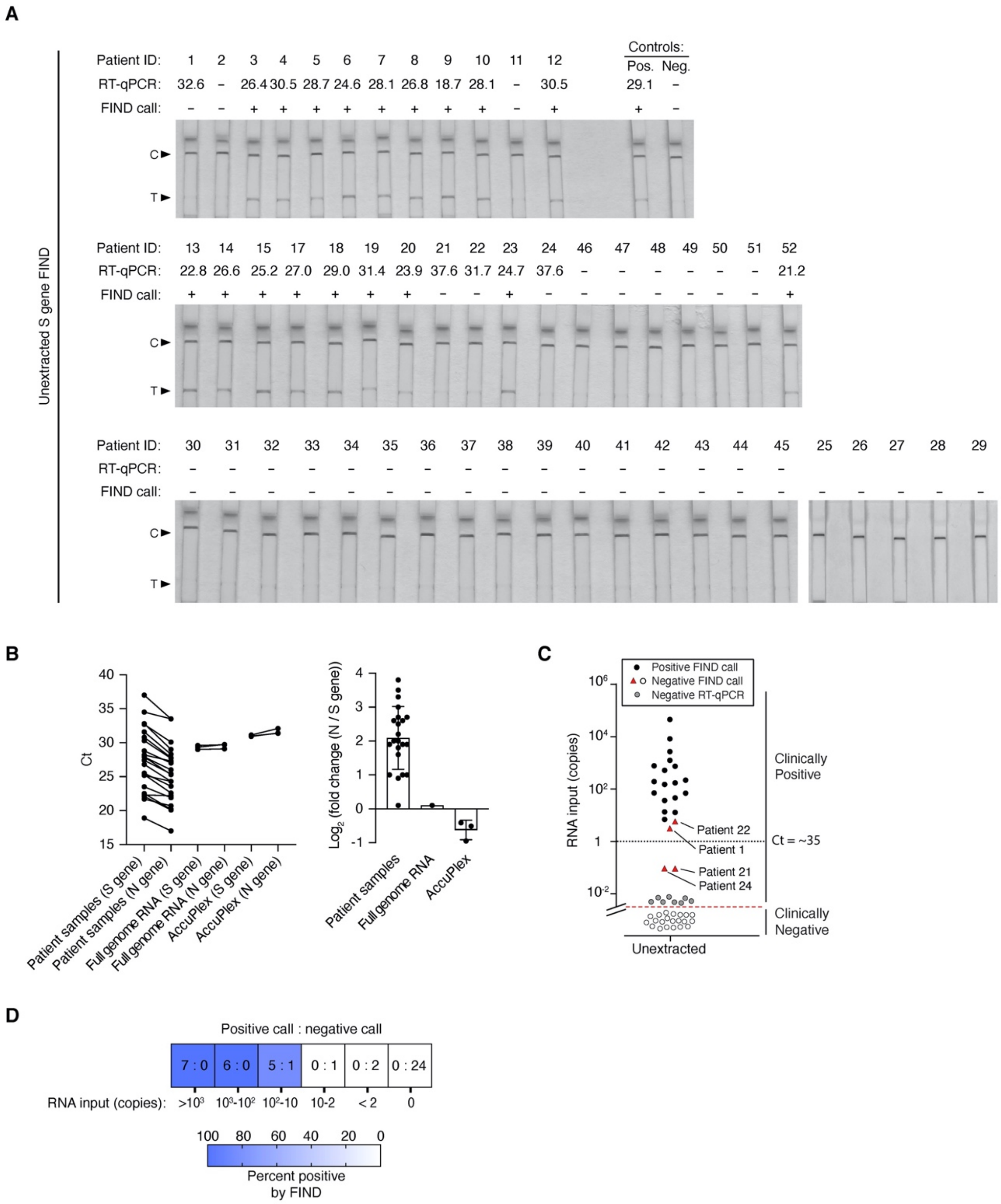


**Figure S5. Detection of SARS-CoV-2 S gene from clinical samples. (A)** Lateral flow strip readouts of S gene FIND performed on patient samples of known infection status. Individual strips are labeled with the test call made within 20 mins of detection (positive (+) or negative (-)). The positive (Pos.) FIND control is 100 copies of synthetic full genome SARS-CoV-2 RNA and the negative (Neg.) FIND control is a water-only input. Negative control samples 25-29 were not screened by RT-qPCR. **(B)** Comparison of Ct values obtained by RT-qPCR targeting SARS-CoV-2 N and S genes on the same input patient samples. In the left panel, matched samples are connected by a solid line. Synthetic full genome SARS-CoV-2 RNA and AccuPlex packaged

SARS-CoV-2 were used as controls as they both contain an equal amount of N and S gene. In the right panel, the difference between the Ct values for each patient sample is plotted and the error bar is +/- 1 standard deviation. **(C-D)** Sensitivity and specificity of S gene FIND on patient samples shown in (A). **(C)** Comparison between S gene SARS-CoV-2 FIND and one-step RT-qPCR performed on the same input samples. The y axis is RNA copies in patient input samples determined by one-step RT-qPCR with standard curve. **(D)** Heatmap displaying the rate of FIND positive tests for detection of SARS-CoV-2 S gene in the 51 patient samples. Values represent the number of positive test calls : number of negative test calls for each condition.

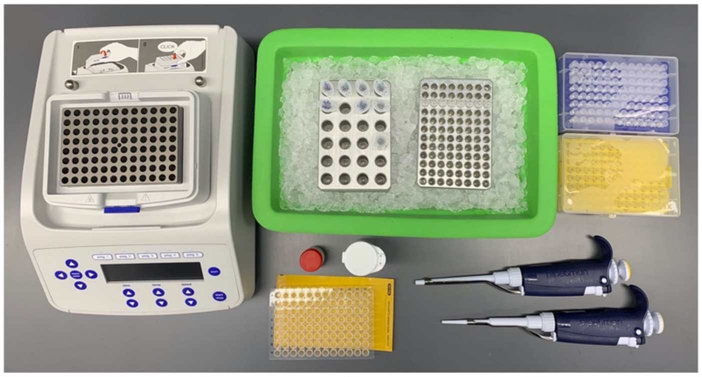

**Figure S6. Equipment required for FIND assay.** FIND only requires a limited set of equipment including micropipettes and disposable plastic tips, a heat block capable of reaching 42°C and 94°C, and plastic microtubes or multi-well plates.

| Oligo name | Alternate name | Oligo sequence | Description of use |
|---|---|---|---|
| CCMS052 | S gene FP1 | GTTTTTCTTGTTTTATTGCCACTAGTCTCTA | SARS-CoV-2 S Gene forward primer |
| CCMS055 | S gene FP2 | TCTTGTTTTATTGCCACTAGTCTCTAGTCAGT | SARS-CoV-2 S Gene forward primer |
| CCMS019 | S gene FP3 | CTCTAGTCAGTGTGTTAATCTTACAACCAGAACT | SARS-CoV-2 S Gene forward primer |
| CCMS053 | S gene FP4 | TCAGTGTGTTAATCTTACAACCAGAACTCAAT | SARS-CoV-2 S Gene forward primer |
| CCMS054 | S gene FP5 | GTGTTAATCTTACAACCAGAACTCAATTACCCC | SARS-CoV-2 S Gene forward primer |
| CCMS067 | S gene RP1 | GAATGTAAAACTGAGGATCTGAAAACTTTG | SARS-CoV-2 S gene reverse primer |
| CCMS014 | S gene RP2 | AGAACAAGTCCTGAGTTGAATGTAAAACTGAGG | SARS-CoV-2 S gene reverse primer |
| CCMS056 | S gene RP3 | AAGAAAGGTAAGAACAAGTCCTGAGTTGAATGT | SARS-CoV-2 S gene reverse primer |
| CCMS030 | S gene RP4 | CCATTGGTCCCAGAGACATGTATAGCATGG | SARS-CoV-2 S gene reverse primer |
| CCMS008 | S gene RP5 | CTCTTAGTACCATTGGTCCCAGAGACATGT | SARS-CoV-2 S gene reverse primer |
| CCMS068 | S gene RP6 | ATCAAACCTCTTAGTACCATTGGTCCCAGA | SARS-CoV-2 S gene reverse primer |
| CCMS073 | | /56-FAM/GAATGTAAAACTGAGGATCTGAAAACTTTG | SARS-CoV-2 S gene reverse primer |
| JQ217 | | CAACTTCCTCAAGGAACAACATTGCCAAAA | SARS-CoV-2 N gene forward primer |
| JQ289 | | GGCTTCTACGCAGAAGGGAGCAGAGGCGGCAG | SARS-CoV-2 N gene forward primer |
| JQ214 | | CAACTGGCAGTAACCAGAATGGAGAACGCA | SARS-CoV-2 N gene forward primer |
| JQ215 | | TCGAGGACAAGGCGTTCCAATTAACACCAA | SARS-CoV-2 N gene forward primer |
| JQ216 | | CCTAGGAACTGGGCCAGAAGCTGGACTTCC | SARS-CoV-2 N gene forward primer |
| JQ218 | | AACTTCTCCTGCTAGAATGGCTGGCAATGG | SARS-CoV-2 N gene forward primer |
| JQ219 | | ATGAAACTCAAGCCTTACCGCAGAGACAGA | SARS-CoV-2 N gene forward primer |
| CCMS041 | | CAATCCTGCTAACAATGCTGCAATCGTGCTA | SARS-CoV-2 N gene forward primer |
| CCMS047 | | CACCAAAAGATCACATTGGCACCCGCAATCC | SARS-CoV-2 N gene forward primer |
| CCMS051 | | CTGAGGGAGCCTTGAATACACCAAAAGATCACA | SARS-CoV-2 N gene forward primer |
| JQ235 | | /56-FAM/TGGAGTTGAATTTCTTGAACTGTTGCGACT | SARS-CoV-2 N gene reverse primer |
| JQ223 | | TGGAGTTGAATTTCTTGAACTGTTGCGACT | SARS-CoV-2 N gene reverse primer |
| JQ220 | | CTTCCTTGCCATGTTGAGTGAGAGCGGTGA | SARS-CoV-2 N gene reverse primer |
| JQ221 | | CTTGGACTGAGATCTTTCATTTTACCGTCA | SARS-CoV-2 N gene reverse primer |
| JQ222 | | GCAGGATTGCGGGTGCCAATGTGATCTTTT | SARS-CoV-2 N gene reverse primer |
| JQ224 | | TGGCCTTGTTGTTGTTGGCCTTTACCAGAC | SARS-CoV-2 N gene reverse primer |
| JQ225 | | TTGAGTCAGCACTGCTCATGGATTGTTGCA | SARS-CoV-2 N gene reverse primer |
| XL12 | | CTCCAGGGAAACTTTGGTGA | MERS N gene RT-qPCR primer |
| XL13 | | CCCATAAAAGCACTGGCTGT | MERS N gene RT-qPCR primer |
| JQ297 | | GGCGCAAGAATTCAGAACCAGAGATA | HCoV-229E N gene RT-qPCR primer |
| JQ298 | | GGATTCCGAGATTGAGATTTGGATTCAC | HCoV-229E N gene RT-qPCR primer |
| JQ303 | | ACTACTCAAGAAGCTATCCCTACTAGG | HCoV-HKU1 N gene RT-qPCR primer |
| JQ304 | | GAGAACGTGAACCTGGTCGACTATTAG | HCoV-HKU1 N gene RT-qPCR primer |
| CCMS083 | | ACGTTGATGTAGGGCCAGAT | MERS S gene RT-qPCR primer |
| CCMS084 | | ACCGTCAGCCTTAGAAACATCA | MERS S gene RT-qPCR primer |
| CCMS086 | | CCGGTGCACCACTTTTGATG | SARS-CoV S gene RT-qPCR primer |
| CCMS087 | | CCCCCTCATAGATGAAGTATGTTGA | SARS-CoV S gene RT-qPCR primer |
| CCMS077 | | CTGTTTGCAACGGCTGTGTT | HCoV-229E S gene RT-qPCR primer |
| CCMS078 | | ACCACCACTCTCAACAGCAA | HCoV-229E S gene RT-qPCR primer |
| CCMS080 | | AAACACCACAGTTCCTCGCA | HCoV-HKU1 S gene RT-qPCR primer |
| CCMS081 | | CCCAAACCATAAGAAACATCCACA | HCoV-HKU1 S gene RT-qPCR primer |
| CC039 | | TCATGGTCCTACATTGTGGAAA | 2009 H1N1 influenza RT-qPCR primer |
| CC040 | | GACACTGAGCTCAATTGCT | 2009 H1N1 influenza RT-qPCR primer |
| GAPDH F | | GGACTCATGACCACAGTCCA | GAPDH RT-qPCR primer |
| GAPDH R | | GATGTTCTGGAGAGCCCCG | GAPDH RT-qPCR primer |
| JQ241 | | /5Biosg/TCAAGCCTCTTCTCGTTCCTCATCACGT | SARS-CoV-2 N gene Hybridization Probe |
| JQ312 | | /5Biosg/CAAGCCTCTTCTCGT | SARS-CoV-2 N gene Hybridization Probe |
| CCMS069 | | /5Biosg/TGCATACACTAATTCTTTCACACGTGGT | SARS-CoV-2 S gene Hybridization Probe |

**Table S1.** List of all primers used in this study.

**Data S1. (separate file)**

Analysis of primer dimers in RT-RPA reactions.

**Data S2. (separate file)**

Bioinformatic analysis of the number of mismatches between FIND assay primers and known variants of SARS-CoV-2. Bioinformatic analysis of the number of mismatches between FIND assay primers and other coronaviruses.

**Data S3. (separate file)**

Data for all patient sample RT-qPCR and FIND assays performed in this study.
